# The developing premature infant gut microbiome is a major factor shaping the microbiome of neonatal intensive care unit rooms

**DOI:** 10.1101/315689

**Authors:** Brandon Brooks, Matthew R. Olm, Brian A. Firek, Robyn Baker, David Geller-McGrath, Sophia R. Reimer, Karina R. Soenjoyo, Jennifer S. Yip, Dylan Dahan, Brian C. Thomas, Michael J. Morowitz, Jillian F. Banfield

## Abstract

1.1

**Background:** The neonatal intensive care unit (NICU) contains a unique cohort of patients with underdeveloped immune systems and nascent microbiome communities. Patients often spend several months in the same room and it has been previously shown that the gut microbiomes of these infants often resemble the microbes found in the NICU. Little is known, however, about the identity, persistence and absolute abundance of NICU room-associated bacteria over long stretches of time. Here we couple droplet digital PCR (ddPCR), 16S rRNA gene surveys, and recently published metagenomics data from infant gut samples to infer the extent to which the NICU microbiome is shaped by its room occupants.

**Results:** Over 2,832 swabs, wipes, and air samples were collected from sixteen private-style NICU rooms housing very low birthweight (<1,500 g), premature (<31 weeks’ gestation) infants. For each infant, room samples were collected daily, Monday through Friday, for one month. The first samples from the first infant and last samples from the last infant were collected 383 days apart. Twenty-two NICU locations spanning room surfaces, hands, electronics, sink basins, and air were collected. Results point to an incredibly simple room community where 5-10 taxa, mostly skin associated, account for over 50% of 16S reads. Biomass estimates reveal 4-5 orders of magnitude difference between the least to the most dense microbial communities, air and sink basins, respectively. Biomass trends from bioaerosol samples and petri dish dust collectors suggest occupancy to be a main driver of suspended biological particles within the NICU. Using a machine learning algorithm to classify the origin of room samples, we show that each room has a unique microbial fingerprint. Several important taxa driving this model were dominant gut colonizers of infants housed within each room.

**Conclusions:** Despite regular cleaning of hospital surfaces, bacterial biomass was detectable at varying densities. A room specific microbiome signature was detected, suggesting microbes seeding NICU surfaces are sourced from reservoirs within the room and that these reservoirs contain actively dividing cells. Collectively, the data suggests that hospitalized infants, in combination with their caregivers, shape the microbiome of NICU rooms.

## 1.3 Background

Hospital acquired infections (HAIs) remain a major problem in the US. One out of every twenty-five patients will experience a HAI, costing the US approximately $30 billion per year [1]. Infants hospitalized in the neonatal intensive care units (NICU) are particularly susceptible to infection due to their underdeveloped immune systems [2, 3]. To protect against infection, infants are often prescribed antibiotics during the first week of life. In fact, antibiotics are three of the six most commonly administered medications in the NICU [4]. This treatment likely kills microbes acquired during the birthing process [5] and promotes a categorically different colonization pattern in preterm infants relative to full term infants [6]. Preterm infants are often colonized by ESKAPE organisms (*Enterococcus spp.*, *Staphylococcus aureus*, *Klebsiella spp.*, *Acinetobacter spp.*, *Pseudomonas aeruginosa*, and other Enterobacteriaceae), which are also the most frequent cause of nosocomial infections [7]. The relatively sterile preterm infant gut microbiome and the high frequency at which infants are colonized by hospital associated microbes, creates a valuable study setting to better understand how the room microbiome is shaped by its occupants. Here, we conducted an experiment to quantify and characterize NICU room microbes to enable comparison with microbiomes that develop in the premature infant gut

The source of early stage gut colonizers in preterm infants has been explored to some extent [8–11]. In a pilot study, we tracked two infants over the first month of life, collecting samples from room surfaces and infant fecal samples [12]. Using an amplicon-EMIRGE approach, which allows for recovery of full-length 16S rRNA genes (~1500 b) [13], as opposed to the more common hypervariable region approach (~150-400 b), we detected the same sequences in room samples before they were detected in gut samples. In a much higher resolution genome-resolved metagenomics study we recently showed evidence for the presence of some infant gut associated strains in the NICU room environment and for exchange of those strains between infant and room environments [24].

Recent genomic studies have shown that the vast majority of strains in the premature infant gut are not shared among infants [5]. Nearly 150 strains were recovered from 10 infants’ fecal samples and only 4 of these were shared. These samples were collected within a month of each other, suggesting that a multitude of strains are available in the NICU at any given point in time, and only a few strains may be widespread, a conclusion supported by the more recent research [Brooks et al. in revision]. However, a few strains were identified in infant fecal samples collected years apart from different infants housed the same NICU [14]. These were referred to as “persister” strains.

A recent study identified 794 antibiotic resistance genes in preterm infant stool samples, 79% which had not previously been classified as associated with resistance [15]. It is possible that these genes provide a competitive advantage for survival in the highly cleaned room environment [16]. However, in our prior work we found that persister strains, which we infer have a room reservoir, were not found to differ significantly in virulence, antibiotic resistance, or metabolism from non-persister strains.

An important question from the perspective of HAI and microbiome establishment of hospitalized premature infants relates to the diversity and biomass distributions over room environments. To address this knowledge gap, we conducted a study with sixteen infants, whose rooms were sampled Monday through Friday from twenty two room locations. We performed droplet digital PCR (ddPCR) on all room samples to directly quantify biomass (2832 samples in total) to determine how biomass varies in the NICU with additional quantification of negative controls. Overall, the findings provide new information about the NICU microbiome and its relationship to room occupant microbiomes.

## 2 Methods

### 2.1 Sample Collection

Infants were enrolled in the study based on the criteria that they were < 33 weeks gestation and were housed in the same physical location within the NICU during the first month of life. Samples were collected Monday through Friday for days of life (DOL) 5-28. Fecal samples were collected from infant diapers and were stored at -20 °C within 10 minutes of collection for short term storage. Shortly after collection, samples were archived and transferred to a -80 °C freezer for long term storage until DNA extraction. All samples were collected after signed guardian consent was obtained, as outlined in our protocol to the ethical research board of the University of Pittsburgh (IRB PRO12100487). This consent included sample collection permissions and consent to publish study findings.

All samples were obtained from a private-style NICU at Magee-Womens Hospital of the University of Pittsburgh Medical Center. Twenty-two of the most frequently touched surfaces were determined by visual observation and health care provider interviews in the weeks leading up to sample collection. Microbial cells were removed from most surfaces using nylon FLOQSwabs (Copan Diagnostics, Brescia, Italy) and a sampling buffer of 0.15 M NaCl and 0.1% Tween20. Samples were collected by one research nurse to ensure consistent sampling technique. Ten square centimeters of each surface was sampled or, for smaller surfaces, the entire surface itself (e.g., isolette knobs and sink basin drain grill). Wipe samples were collected from the floor and exterior top of the isolette using Texwipe TX1086 wipes (Texwipe, Kernersville, NC, USA). Before collecting each wipe sample, the collector would put on latex examination gloves and clean these gloves with an isopropanol wipe. The wiped surface area was approximately forty-eight square centimeters or, for smaller surfaces, the entire surface itself (e.g., isolette top). A wipe was also used to collect microbial cells at the exterior facet of the heating, ventilation and air conditioning (HVAC) system. The wipe was suspended via airflow on the exterior (upstream) face of the MERVE 8 pleated filter, the zone in which supply and return air are mixed before thermal and humidity treatment of the airstream for four days. Features of the HVAC system are described in detail in a recently published paper [18].

Air samples were collected using the NIOSH two-stage bioaerosol cyclone 251 sampler [19] and a suspended petri dish method [20]. The NIOSH sampler collected samples continuously Monday through Friday, comprising approximately 96 hours of sampling at 3.5 L/minute (total volume sampled = 20 m3). Petri dish samples were suspended approximately one meter below the drop ceiling in the corner of the room that was the furthest away from the sink. These samplers were maintained in place for the duration of the infant’s stay. Petri dish “cooler” samples are plates that were taped to the top of a cooler which collected abiotic aerosol data [18]. At the end of the sample collection period, all samples were placed in a sterile transport tube and stored within 10 minutes at -80 °C until further processing.

### 2.2 DNA extraction and PCR amplification

DNA was extracted using either the MO BIO PowerSoil DNA Isolation kit (single tube extractions) or PowerSoil-htp 96 Well DNA Isolation kit (MoBio Laboratories, Carlsbad, CA, USA). For DNA extracted from feces with the 96-well kit fecal samples were kept frozen on dry ice and added to individual wells of the bead plate and stored at -80°C until extraction. The day of extraction Bead Solution and Solution C1 were added and the plates were incubated at 65°C for 10 minutes. The plates were shaken on a Retsch Oscillating Mill MM400 with 96-well plate adaptors for 10 minutes at speed 20. The plates were rotated 180° and shaken again for 10 minutes at speed 20. All remaining steps followed the manufacturer’s centrifugation protocol. For swab samples the heads were snapped at the perforation into the wells of the bead plate and stored at - 80°C. The day of extraction the Bead Solution and Solution C1 were added and the plates were incubated at 65°C for 10 minutes. The plates were shaken on a Retsch Oscillating Mill MM400 with 96 well plate adaptors for 5 minutes at speed 20. The plates were rotated 180° and shaken again for 5 minutes at speed 20. The Solution C2 and C3 steps were combined (200 μl of each added) to improve DNA yield. All remaining steps followed the manufacturer’s centrifugation protocol.

Wipe samples were stored in a sterile 250 mL tissue culture flask upon collection and thawed on ice before extraction. Cells were dislodged from wipes in a protocol adapted from Yamamoto *et al*. [17]. Briefly, 150 mL of dislodging buffer was poured into a flask (1X PBS, 0.04% Tween 80, passed through a 0.2 μm filter) and the flask was shaken vigorously for one minute. Supernatant was then decanted into a 250 mL disposable filter funnel with a pore size of 0.2 μm (Thermo Scientific, Waltham, MA, USA) and the filter was then placed in a MoBio PowerWater extraction tube. PowerWater extraction followed manufacturer recommendations.

Droplet digital PCR (ddPCR) was adapted from a method previously published on quantification of 16S rRNA templates in infant fecal samples [5]. The only deviation from the previous method was that a diluted gDNA template of 1:10 instead of 1:1000 was utilized. Both MiSeq library preparation and ddPCR were performed in 96-well plate format. Each plate had three no template PCR controls, one no template extraction control, and three positive controls containing varying concentrations of purified *E. coli* gDNA. Counts from the negative control types were averaged across type and the highest was used to correct for contaminant counts in sample data.

### 2.3 Sequencing preparation and sequencing

Genomic DNA from room samples were subjected to 16S rRNA V3-4 MiSeq library preparation which included dual-barcoded multiplexing with a heterogeneity spacer for higher sequence quality [22]. Two microliters of 5X concentrated gDNA template was used in the reaction and run at 35 cycles. Amplicons were purified using the Just-a-Plate PCR normalization and purification kit (Charm Biotech, San Diego, CA, USA). Equal amounts of each sample were sent to the University of California Davis DNA Technologies Core Facility (http://dnatech.genomecenter.ucdavis.edu) and run on a MiSeq with v3 300PE chemistry.

Illumina library construction for infant fecal samples followed standard protocols at University of California QB3 Vincent J. Coates Genomics Sequencing Core Facility (http://qb3.berkeley.edu/gsl/). Briefly, gDNA was sheared using a Covaris to approximately 600 bp and 1000 bp. Wafergen’s PrepX DNA library prep kits were used in conjunction with the Apollo324 robot following factory recommendations (Integenx). Thirteen cycles of PCR were used during library construction. Libraries were added at 12 samples per lane, in equimolar amounts, to the Illumina HiSeq 2500 platform. Paired-end sequences were obtained with 150 cycles and the data processed with Casava version 1.8.2. Raw read data were deposited in the NCBI Short Read Archive (Bioproject PRJNA376566, SRA SUB2433287).

### 2.4 16S amplicon data processing

The LotuS 1.562 pipeline in short amplicon mode was used for quality filtering, demultiplexing, and OTU picking [23]. LotuS was run with the following command line options: ‘-refDB SLV,GG -highmem 1 -p miseq -keepUnclassified 1 -simBasedTaxo lambda -threads 10.’ The OTU data was rarefied to 1,000 sequences per sample, without replacement, unless explicitly stated. OTU table and LotuS log files are available in Additional file 1.

### 2.5 Metagenomic data from infant gut samples

For comparative purposes, this study made use of previously published infant metagenomic data from 290 fecal samples collected from infants housed in the NICU rooms studied here (~800 Gb of 150 bp paired-end reads). Methods for data analysis are described within this publication [24].

## 3 Results

### 3.1 Sequencing summary and contamination removal

In total, 2832 room samples were processed through a MiSeq library preparation protocol. After quality filtering and demultiplexing, 84,939,529 read pairs were generated. These reads were clustered into 18,093 OTUs. Using a ratio OTU (ROTU) method that leverages biomass quantification and sequencing of negative controls [25], 269 OTUs and 925 samples were removed from the dataset when using an ROTU threshold of 0.001. A second *in silico* contamination cleaning method was applied [26], which removed an additional 323 OTUs and 1 sample. In total, approximately 3% of generated OTUs and 33% of samples present too weak of a signal to confidently distinguish them from negative control signatures.

### 3.2 Biomass and taxonomic variation across petri dish replicates

Biological and technical replicates performed for petri dish plates established the reproducibility of extraction of DNA from petri dish swabs and provided evidence for highly reproducible ddPCR measurements (Additional file 2). The highest standard deviation in ddPCR values for biological replicates in a single room was 106,760 copies/sample (infant 6’s petri plates; mean = 99,677) and for technical replicates, the largest standard deviation was 15,534 copies/sample (infant 12’s petri plates, mean = 81,044). The lowest standard deviation for biological replicates was 1,981 copies/sample (infant 1’s petri plates, mean = 13,785) and 737 copies/sample for technical replicates (infant 11’s petri plates, mean = 32,396). Overall, this equates to a reproducibility range of 2.69 to 6.87 × more reproducibility across technical ddPCR runs relative to biological replicates, with an average reproducibility ratio of 5.37 × better for technical replicates.

### 3.3 Biomass varies significantly across sample type

16S rRNA gene copies were quantified for 2,883 samples using ddPCR and showed day-to-day variation ranging from approximately 4 to 33000 16S rRNA copies/cm2 (Figure 1a). Samples from the HVAC system had the highest biomass of all types and bioaerosol samples had the lowest (Additional file 3 a and b). Sinks had the highest biomass of the swabbed samples and hands had the lowest average median template count (Figure 1b). Petri dishes suspended from the ceiling had the lowest biomass relative to other passive dust collectors, whereas the nurse’s station dishes contained the highest bacterial load. The infant room consistently had higher template counts than the hallway bioaerosol samples. Overall, the median biomass varied over 4 orders of magnitude across all sample types.

**Figure 1:**
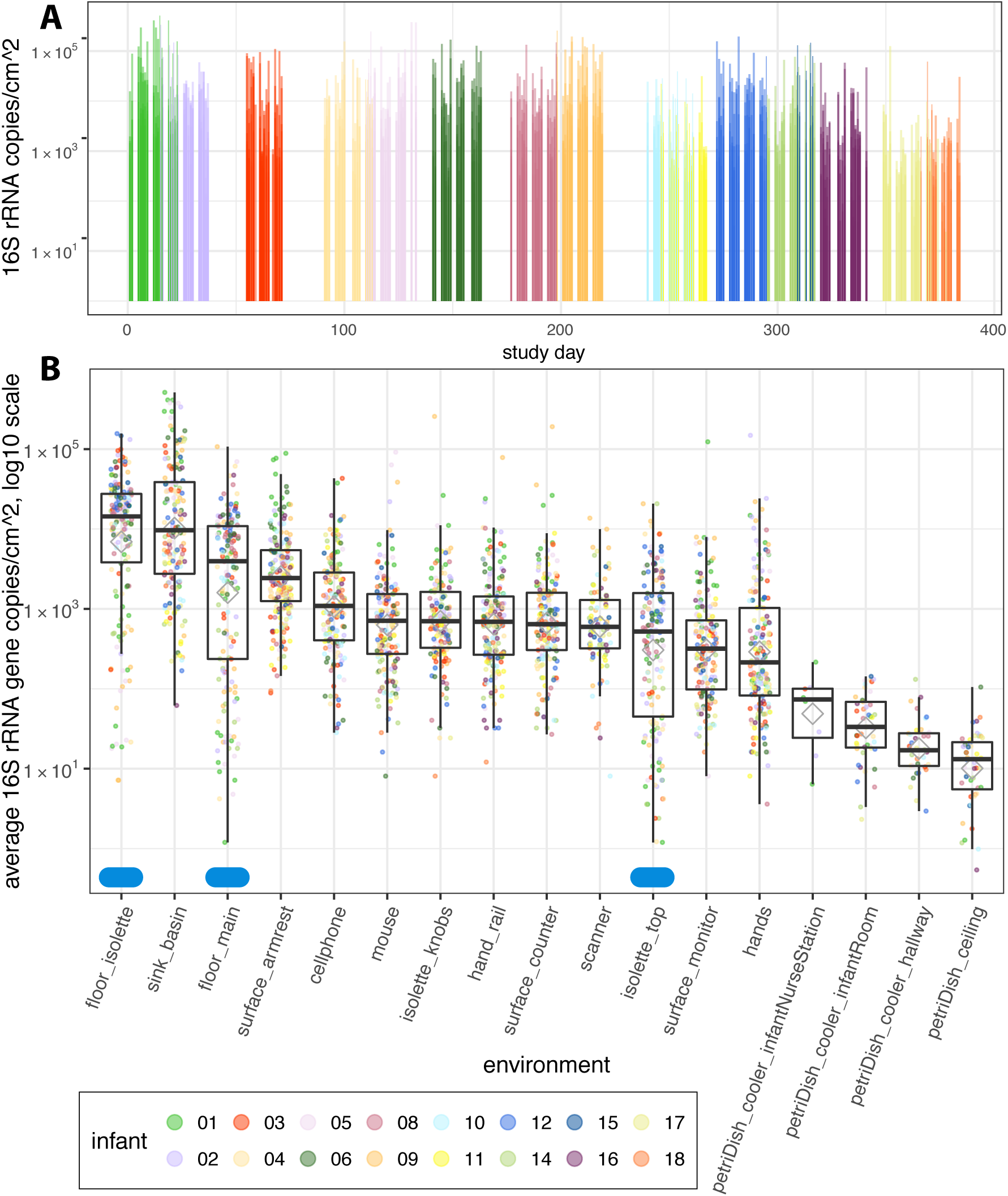
Biomass varies by 4-5 orders of magnitude in a NICU. 16S rRNA template copy number was quantified via ddPCR. (A) Biomass was averaged across all swab and wipe samples for each sampling day and plotted on a timeline to visualize variation in biomass over the sampling campaign. (B) Each dot reflects the average across triplicate runs. Grey diamonds represent averages per environment. Blue ellipses along the x-axis represent samples collected using a wipe method. All other samples were collected with swabs or using a petri plate to collect settled dust (noted in label). All counts are normalized to represent one day of collection.

### 3.4 Skin associated taxa dominate the NICU surface environment

The microbial communities in most NICU environments were highly uneven and were dominated by 5-10 OTUs (Figure 2). 41% and 55% of all amplicon reads belong to the top five and ten OTUs in the NICU, respectively (Figure 2 and Table 1). Most of these taxa are human associated with many commonly associated with the skin (*Corynebacterium*), mouth (*Streptococcus*), or nose (*Staphylococcus*). SourceTracker v1.0.1 [27] was run using skin, oral, and fecal samples from the American Gut project as the putative source database with NICU samples labeled as “sink” samples. Skin was the most likely contributor to taxa in the NICU, accounting for upwards of 50% of the most probable sources, followed by oral and fecal samples (Additional file 4).

**Figure 2:**
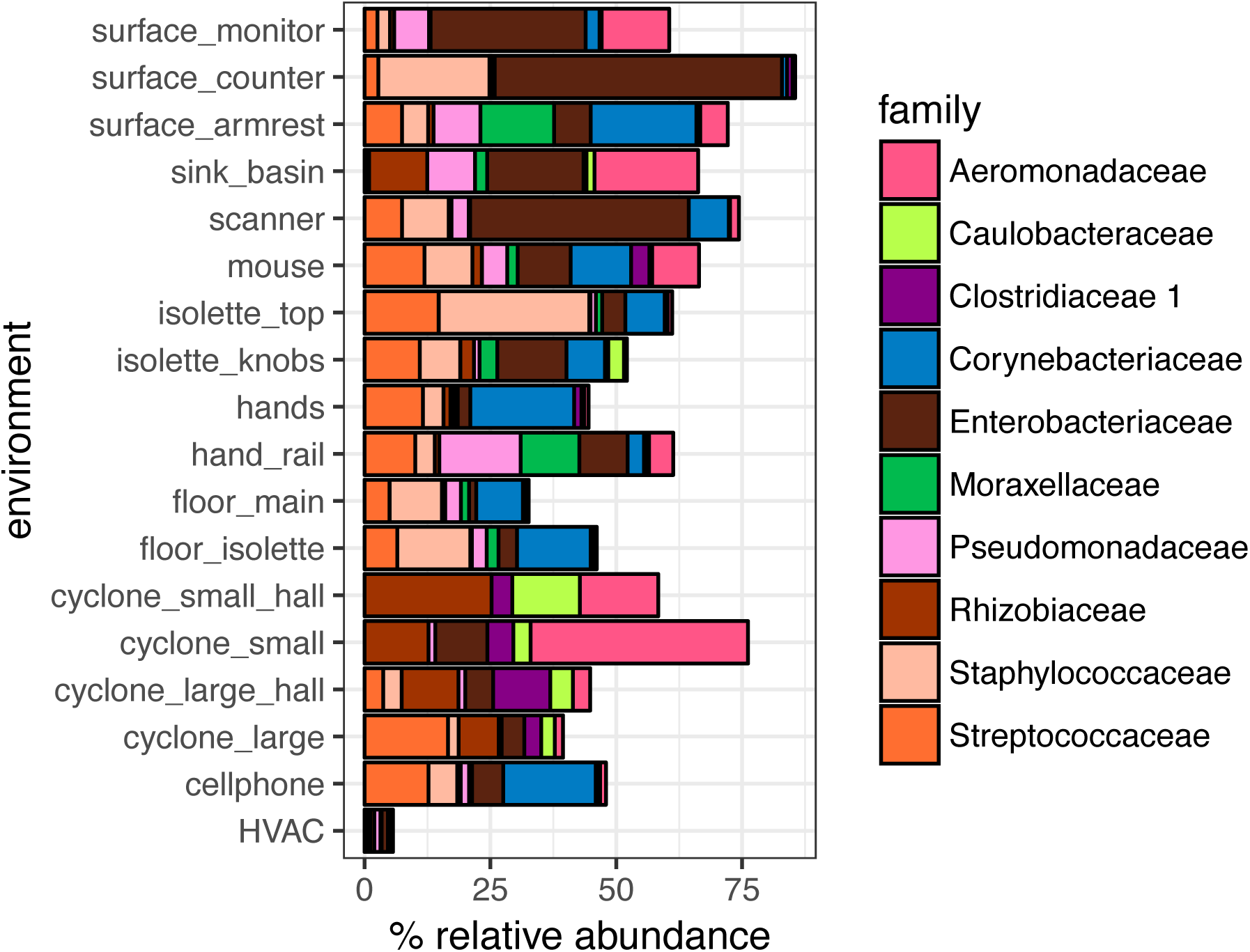
Top 10 NICU OTUs comprise > 50% of NICU taxa. Amplicon data from a 16S rRNA V3-4 workflow is plotted for each environment. Only the top 10 OTUs, determined from averages across all samples, are plotted. Each OTU is colored by its family-level classification.

**Table 1:** Top 10 OTUs in the NICU.

| OTU | Kingdom | Phylum | Class | Order | Family | Genus | Species | % Abundance |
| --- | --- | --- | --- | --- | --- | --- | --- | --- |
| OTU_5 | Bacteria | Proteobacteria | Gammaproteobacteria | Enterobacteriales | Enterobacteriaceae | Klebsiella | ? | 12.9 |
| OTU_6 | Bacteria | Firmicutes | Bacilli | Bacillales | Staphylococcaceae | Staphylococcus | ? | 7.3 |
| OTU_4 | Bacteria | Actinobacteria | Actinobacteria | Corynebacteriales | Corynebacteriaceae | Corynebacterium | ? | 7.1 |
| OTU_7 | Bacteria | Firmicutes | Bacilli | Lactobacillales | Streptococcaceae | Streptococcus | ? | 6.9 |
| OTU_9 | Bacteria | Proteobacteria | Gammaproteobacteria | Aeromonadales | Aeromonadaceae | Aeromonas | ? | 6.9 |
| OTU_10 | Bacteria | Proteobacteria | Alphaproteobacteria | Rhizobiales | Rhizobiaceae | Rhizobium | ? | 4.5 |
| OTU_8 | Bacteria | Proteobacteria | Gammaproteobacteria | Pseudomonadales | Pseudomonadaceae | Pseudomonas | ? | 3.7 |
| OTU_11 | Bacteria | Proteobacteria | Gammaproteobacteria | Pseudomonadales | Moraxellaceae | Acinetobacter | ? | 2.3 |
| OTU_30 | Bacteria | Firmicutes | Clostridia | Clostridiales | Clostridiaceae 1 | Clostridium<br>sensu stricto 1 | ? | 1.9 |
| OTU_32 | Bacteria | Proteobacteria | Alphaproteobacteria | Caulobacterales | Caulobacteraceae | Brevundimonas | ? | 1.8 |

Samples collected from the HVAC system had the highest bacterial diversity with 405 OTUs on average per sample, whereas bioaerosol samples had the lowest, with 13 (Additional file 5a). The HVAC samples had the highest Shannon community evenness, followed by floor wipes, and the bioaerosol samples had the lowest Shannon diversity (Additional file 5b). Thus, overall, the HVAC had highly even consortia with high diversity. This is expected due to the way that the HVAC sample was collected, with metric tons of air passing through the collection wipe before sequencing [18]. The NICU room air was also found to have low biomass and low diversity, with strong dominance by members of the *Aeromonadaceae* in the small size fraction and *Streptococcaceae*, *Rhizobiaceae*, *Clostridiaceae* in the large size fraction.

All touched surfaces had similar numbers of OTUs per sample, although the surface monitors showed the most unevenness (Additional file 5). These surfaces were dominated by similar groups of microbes. Although many touched surfaces were associated with skin-associated bacteria, gut associated Enterobacteriaceae OTUs also dominated environments such as the surface monitors, counter tops, and scanners (Figure 2). In contrast, the sink basins had comparatively low numbers of OTUs per sample (Additional file 5a), in part due to the high dominance by four bacterial groups (Figure 2).

### 3.5 Biomass suggests growth patterns in sink basins

A range of 29 to 38 sink basin samples per weekday were collected from 14 unique sink basins. When comparing biomass trends across days (Figure 3a), a distinct pattern of decreasing biomass is apparent in sink samples relative to other swabbed environments. In comparing Shannon diversity across weekdays (Figure 3b), bacterial diversity in Tuesday versus Friday samples were the most distinct, whereas biomass was most different in Monday versus Thursday samples (Wilcoxon rank sum, Bonferroni adjusted *p* = 0.47 and 0.012, respectively). Sink basins were cleaned approximately every twenty-four hours, but less frequently on the weekends, so the elevated biomass at the beginning of the week may be due to regrowth of sink adapted taxa throughout the weekend (e.g., *Rhizobiaceae*, *Pseudomonas*, *Aeromonas*, and *Enterobacteriaceae*). The increase in Shannon diversity from Monday to Friday strengthens this inference.

**Figure 3:**
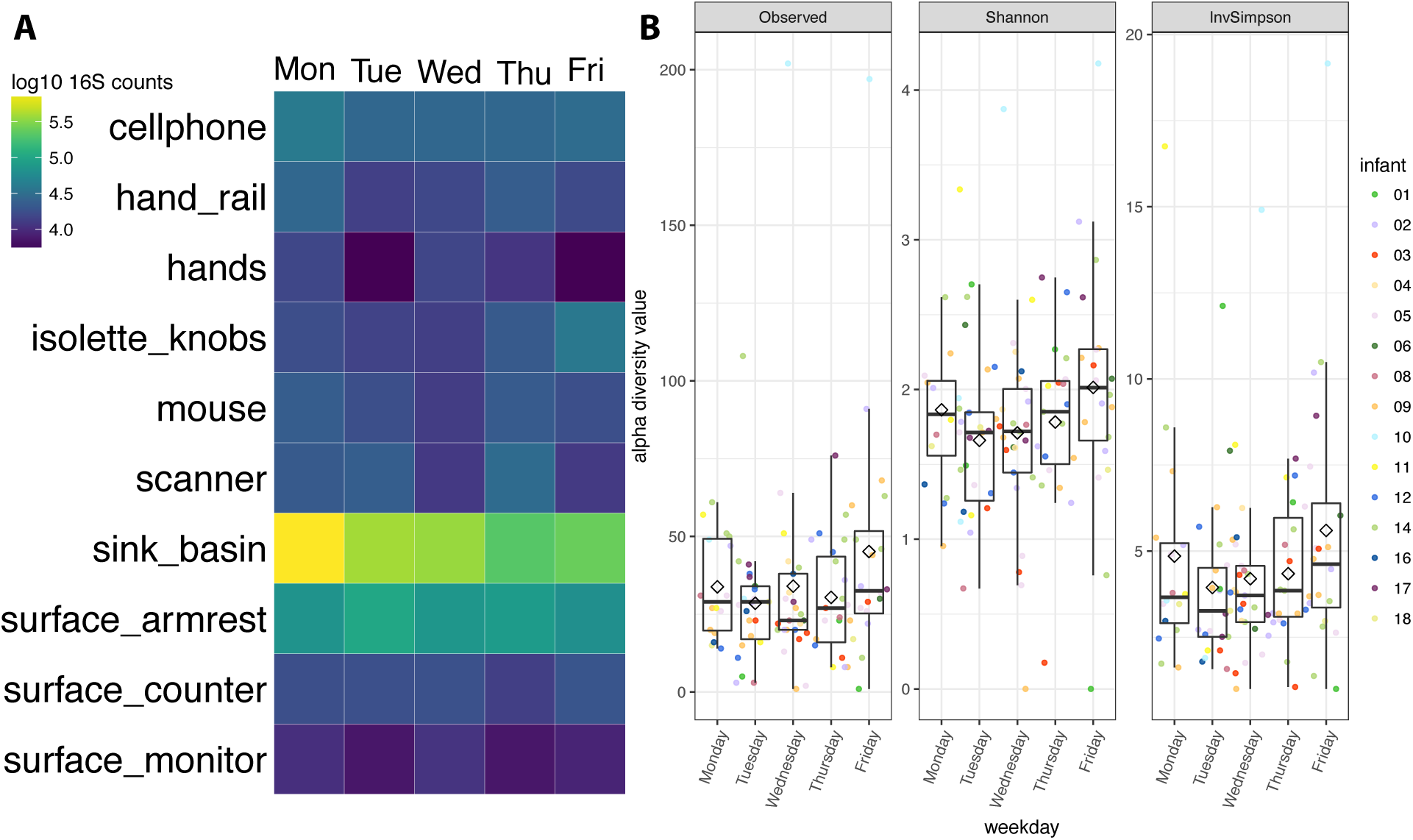
Growth detected in NICU sink samples. 16S rRNA template copy number was quantified via ddPCR. Average copy number was averaged for each weekday and swabbed environment and displayed in this heatmap (a). 16S rRNA amplicon data was used to calculate number of OTUs, Shannon, and Inverse Simpson diversity metrics for sink basin samples (b). Black diamonds represent averages per weekday.

### 3.6 NICU rooms harbor a unique microbial signature

Using a support vector machine (SVM) classifier with a linear kernel [28], we determined that each room’s microbiome contained a unique microbial fingerprint. We could predict the room origins with an overall accuracy of 56% (when we knew the room’s origin but withheld that information from the classifier), which is 5x better than random chance (Figure 4). The use of ROTU over a standard pipeline achieved an increase in accuracy of approximately 16%. Typically, the most confusion occurred between samples that were collected at similar times, although infants that had similar gut communities had decreased prediction accuracy (e.g. infants 2, 3, and 8). Important OTUs driving the SVM model are plotted and listed in Additional file 6 and Table 2. Interestingly, there is an overlap between room specific OTUs that drive the SVM model and occurrence of these taxa in the gut of infant occupants. For example, the most visible signature in SVM taxa comes from a spike in *Veillonella* in infant 6’s room on DOL 18 (Additional file 6). A major increase of *Veillonella* in infant 6’s gut occurred on DOL 16 (ref http://ggkbase.berkeley.edu/project_groups/human-gut-metagenome-sloan-infants and Additional file 7). The same pattern is seen for infant 8, and in fact, most infants that contain *Veillonella* have strong SVM signals associated with their room. The second strongest signal from the SVM model comes from a *Clostridium* OTU. This group is present in infants 2, 3, and 8’s room samples and it strongly contributes to the SVM model prediction. All three of these infants have high abundances of *Clostridium*.

**Figure 4:**
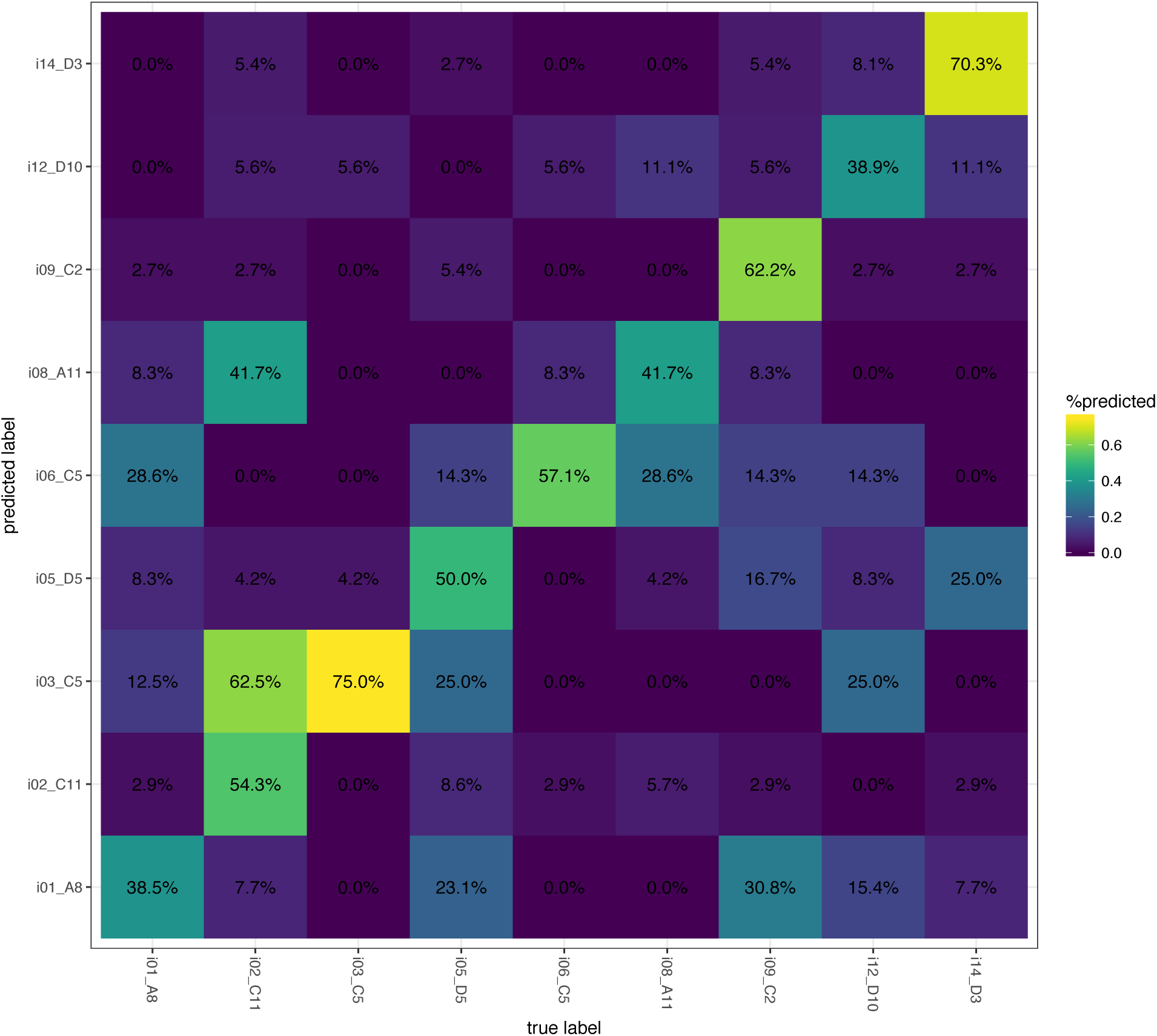
NICU rooms have a unique microbial signature. 16S rRNA amplicon data was split into training, test, and validation sets to train, test, and validate a support vector machine classifier. The confusion matrix plots the accuracy of our model on the validation dataset. Percentages note the number of times a sample was predicted to belong to a room-infant pairing divided the total number of samples for that room-infant pairing. The heat coloring is based on shown percentages.

**Table 2:** Most important variables to SVM model.

| OTU | Kingdom | Phylum | Class | Order | Family | Genus | Species |
| --- | --- | --- | --- | --- | --- | --- | --- |
| OTU_29 | Bacteria | Firmicutes | Clostridia | Clostridiales | Clostridiaceae 1 | Clostridium sunsu<br>stricto 1 | uncultured organism |
| OTU_39 | Bacteria | Actinobacteria | Actinobacteria | Micrococcales | Micrococcaceae | Rothia | uncultured organism |
| OTU_41 | Bacteria | Firmicutes | Bacilli | Bacillales | Family XI | Gemella | ? |
| OTU_30 | Bacteria | Actinobacteria | Actinobacteria | Micrococcales | Micrococcaceae | Kocuria | ? |
| OTU_45 | Bacteria | Actinobacteria | Actinobacteria | Actinomycetales | Actinomycetaceae | Actinomyces | ? |
| OTU_43 | Bacteria | Firmicutes | Bacilli | Bacillales | Alicyclobacillacea<br>e | Tumebacillus | uncultured Firmicutes bacterium |
| OTU_76 | Bacteria | Firmicutes | Clostridia | Clostridiales | Family XI | Peptoniphilus | ? |
| OTU_74 | Bacteria | Actinobacteria | Actinobacteria | Actinomycetales | Actinomycetaceae | Actinomyces | uncultured organism |
| OTU_28 | Bacteria | Firmicutes | Negativicutes | Selenomonadales | Veillonellaceae | Veillonella | uncultured organism |
| OTU_66 | Bacteria | Firmicutes | Bacilli | Lactobacillales | Streptococcaceae | Streptococcus | ? |

### 3.7 Composition of persister taxa in the room echoes infant gut composition

To visualize the distribution of families with representative strains known to persist in infants over multi-year periods [14, 15], we collapsed each study day and infant pairing by averaging all amplicon abundance data across environments (Additional file 8, “average” panel). In this analysis, the subset of all OTUs that belonged to a persister family was assigned a distinct color but often one OTU could be distinguished within a family. However, due to high abundance, we gave OTU_5 (an *Enterobacteriaceae*) dedicated coloring. Surprisingly, persister families account for > 50% of the data at many time points.

Episodes of particularly high persister family abundance occurred in rooms housing infants 1, 9, 12, and 16. To better visualize which samples contributed to the averaged data (Additional file 8, “average” panel), we also plotted data for the specific environments for which we had the most samples (armrests and sinks). Both the armrests and sinks are dominated by these groups of organisms during these episodes, but *Staphylococcaceae* OTUs are much more abundant in armrest samples relative to sinks. Two dominant Pseudomonas OTUs that comprised 70% and 24% of all Pseudomonadaceae (OTU_8 and OTU_15, respectively) were detected throughout the time series, but were at very low abundance in armrest samples over long time spans.

#### Composition of persister taxa in infant 9

Since the room data for infant 9 had a strong signal for persister groups, we analyzed samples from all environments separately to visualize temporal patterns (Figure 5a). Persister groups dominated most of infant 9’s room samples, with cellphones having the fewest and scanner and surface counter samples having the most persister groups per sample. The red lines in Figure 5a highlight the time point where a major increase in relative abundance of *Enterobacteriaceae* taxa occurred in infant 9’s gut (Figure 5b and Additional file 7). This group is present in multiple room environments prior to the increase, particularly associated with the isolette and armrest. At subsequent time points, this group becomes highly prominent in some room environments (e.g., scanner and surface counter).

**Figure 5:**
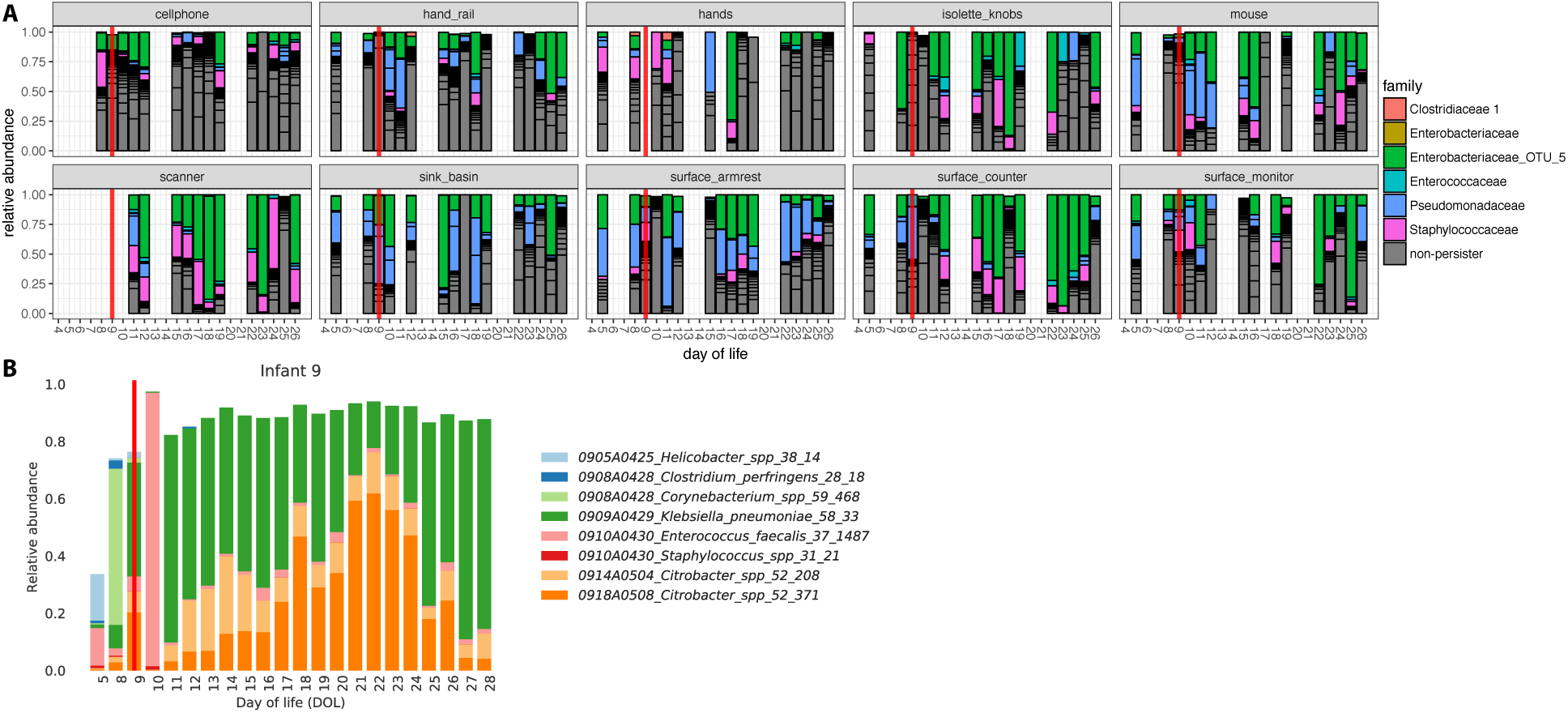
‘Persister taxa in the room reflect composition of the infant gut. Infant 9’s room amplicons are plotted for each swabbed environment (a). Colored are OTUs that belong to a persister lineage. Red lines highlight day of life 9, which coincides with an increase of several *Enterobacteriaceae* taxa in the infant gut (b). (b) is the microbial profile for fecal samples generated via genomes recovered from a metagenomics approach.

OTUs belonging to the persister groups cannot be confidently classified genus level via 16S rRNA gene sequencing [29], and since *Enterobacteriaceae* dominates the gut of infant 9, we leveraged room and fecal sample context to infer a possible identity for OTU_5. Using OTU_5’s reference sequence as a query, we ran ublast [30] on a database of 16S rRNA genes reassembled from infant 9’s fecal metagenomic samples using the REAGO algorithm (Yuan *et al.*, 2015). The top hit to our 429 bp query was 99.5% identical (2 mismatches) and came from several of infant 9’s fecal samples. Most of the top hits have the entire 16S rRNA gene recovered from the REAGO assembly (~1,520 bp). These fecal sequences were searched against the Silva database (SLV_119_SSU) and returned identical, full-length matches to *Klebsiella pneumoniae*. While this is an extrapolation from the V3-4 region, it is possible that OTU_5 in the room is a *Klebsiella* and may be *Klebsiella pneumoniae*, the dominant bacterium colonizing infant 9.

## 4 Discussion

The first question that we aimed to answer in this study related to how biomass varies across a NICU. Using ddPCR to quantify 16S rRNA gene copy number, we show biomass density varies across NICU surfaces by 4-5 orders of magnitude (Figure 1). Surprisingly, the floor in front of the infant’s isolette had the highest density of microbes relative to any other environment within the NICU. Naively, it may seem intuitive that the region with the most foot traffic, e.g. the floor at the main entrance of the NICU, would have the highest biomass. While the main entrance floor has a high density, it is significantly lower than the floor in front of the isolette. This finding may be due to the increased occupancy at the isolette versus the main entrance, where occupancy is more transient.

Petri dish data also suggest that higher levels of human activity drive higher amounts of microbial deposition in the room environment. The nursing station has higher petri dish-associated biomass than the infant room, followed by the hallway (Figure 1). This outcome occurred despite the fact that the infant room and hallway coolers collected dust at the same height (1 m), whereas the nurse station collector was at approximately double the height (1.8 m). As height above the floor increases, detection of resuspended particles from dust decreases exponentially [32, 33]. This finding suggests that floor dust is not the main source of biological particles accumulated in the petri dishes, but rather the microbes are human-derived. Greater occupancy or rigor of activity [34] at the nursing station compared to the infant room and hallway likely explains this result.

A recently published study noted a stronger occupancy signal from the occupancy sensors in the infant room compared to the hallway [18]. The occupancy signal directly overlapped with the coarse particle signal (which detected particles > 10 μm in diameter). This signal was interpreted to indicate that resuspension or deposition of particles from occupants is the largest contributor of aerosolized particles in the NICU. In the current study, our Petri dish ceiling analyses suggest a similar conclusion for settled particles, but in this case based on biological data.

If occupancy is a key feature of the NICU environment, one would expect human associated microbes to dominate in most room environments. We found that 5-10 OTUs account for most of the amplicon data and a majority of these are typically skin, nose, or fecal associated (Figure 2). The enrichment of human associated taxa is likely due to tight control of the building envelope via HVAC treatment [35] combined with a strict cleaning schedule.

An interesting finding of this study related to the change in biomass and microbial community structure of the sink basins over the course of the week. We attribute this pattern to the room cleaning regime, which is more limited on weekend days than during the week. On Mondays, the sink biomass is highest (Figure 3a) and communities are relatively uneven (Figure 3b), presumably due to extensive growth of a few sink-associated taxa over the weekend. More intensive cleaning of the sink early in the week likely removes the majority of biomass, which is comprised of the sink-adapted taxa and enables detection of a wider diversity of low abundance, poorly adapted or transient, taxa.

The second question addressed in our study related to the taxa that dominate NICU surfaces. To investigate this, it was necessary to adapt a method to eliminate spurious contaminant-based signals in data from low biomass samples [25]. The ROTU cleaning method implemented here to clean data of spurious OTUs and contaminants *in silico* was made possible due to the availability of ddPCR quantification of negative controls. This capability is particularly important for NICU studies since the rooms are cleaned regularly, causing low biomass levels to be present in many samples. Some of the bacteria that we conclude were introduced in sample processing are skin associated, although many types of taxa were encountered. After accounting for contamination, we conclude that human associated taxa dominate most surfaces.

Human associated taxa are likely sourced and trafficked throughout the NICU by healthcare providers [36] and many hand hygiene studies have reported as much [37]. Here, we implemented a machine learning classifier to address the possibility that infants and their caretakers shape the microbiome to be distinctive in each room. Our model reliably classified samples of unknown origin to their correct room-infant pair at an accuracy two times better than a recently published office microbiome study [28] and achieved predictive power five times better than random chance. This outcome suggests that NICU rooms are more personalized than other common built environments. There are typically a larger variety of activities and people in office spaces and air treatment is less (lower air exchange rates and less filtration). The combination of less frequent cleaning, increased occupancy, and more unfiltered outdoor air supply drives many of the differences between other common indoor environments and the NICU. The more unique room signal based on NICU room microbes suggests a localized source of bacteria, since a more diffuse source would lower prediction accuracy. A similar result was recently described in a microbiome study conducted in a Chicago hospital [38]. Microbial community similarity increased between patients’ hand and floor samples over time, highlighting the exchange between patient and room. Interestingly, infants in this cohort are rarely removed from their isolettes, so room specific microbiomes were likely mediated by health care providers, rather than direct infant interaction with surrounding room surfaces.

Finally, we tested for patterns of association between room occupants and NICU room environments. We found that many taxa driving our machine learning model for the room microbiome were from groups also present in the gut of the infant occupant. Other signals came from *Firmicutes* and *Actinobacteria* not affiliated with the infant gut and that were relatively uniquely detected in certain rooms. Focusing on the subset of taxa that are gut colonizers, we show a relatively high abundance of these taxa throughout the sampling campaign (Additional file 8). Episodes where persistent families increase and 2-3 OTUs comprise > 30% of the data across all environments occurred several times throughout the study (*e.g.*, in infants 9, 12, and 16). These OTUs are detected in low abundance in the room before detection in the gut (Figure 5). Once in the infant gut, a far more favorable environment for growth and reproduction than on exposed hospital surfaces, bacterial density can reach nearly 10 billion cells per gram [5]. After a spike in relative abundance in the gut, we see these organisms increase in abundance in the room environment. It is impossible to resolve room 16S rRNA amplicon data to the strain-level in order to make claims that the same gut bloom resulted in a subsequent expanded appearance in the room. Potentially, infant 9’s dominant gut *Klebsiella pneumonia* may be linked to an increased abundance of Enterobacteriacaea in the room. Interestingly, the same strain of *K. pneumoniae* found in the gut was detected years apart in different infants within this NICU [14].

## 5 Conclusions

Based on the current study, we conclude that two factors shape room microbiomes. First, our taxa identifications and occupancy results extend prior findings of a strong link between human activity levels and room microbiology [12, 18, 24, 34] In fact, this connection appears to be strong enough to give rise to a relatively unique room microbiome character. Second, environmental stresses, likely associated with cleaning [12, 16, 39–41], likely selectively shape NICU microbiomes, primarily by selecting for microbial specialists that can both thrive in the gut and tolerate the NICU environment. While daily cleaning substantially lowers the bioburden in the NICU [42], the harshest cleaning methods cannot sterilize hospital surfaces [7]. Creative new approaches to displace or prevent entrenchment of these NICU specialists, possibly through prebiotic building materials or clever probiotics, may present opportunities to break the room-occupant cycle.

## 6 Declarations

## 7 Ethics approval and consent to participate

All samples were collected after signed guardian consent was obtained, as outlined in our protocol to the ethical research board of the University of Pittsburgh (IRB PRO12100487). This consent included sample collection permissions and consent to publish study findings.

## 8 Consent for publication

Consent was obtained to publish study findings (IRB PRO12100487).

## 9 Availability of data and materials

Raw read data were deposited in the NCBI Short Read Archive (Bioproject PRJNA376566, SRA SUB2433287). OTU table and LotuS log files are available at https://goo.gl/zQf7FY.

## 12 Authors’ Contributions

JFB, MJM, and BB conceived of the project. RB organized cohort recruitment and sample collections. BAF conducted nucleic acid extractions. BB, DG, SRR, KRS, and DD conducted ddPCR quantifications and MiSeq library preparations. BB conducted the metagenomic assemblies, BCT provided bioinformatics support and MRO contributed to data analysis. BB and JFB wrote the final manuscript. All authors have read and approved the manuscript.

## 13 Acknowledgments

**Additional file legends**

Additional file 1: **OTU table and LotuS log files.** Output from the LotuS pipeline is provided including raw OTU table, accompanying mapping file with cohort and ddPCR count data, and accompanying log files.

**Additional file 2:**
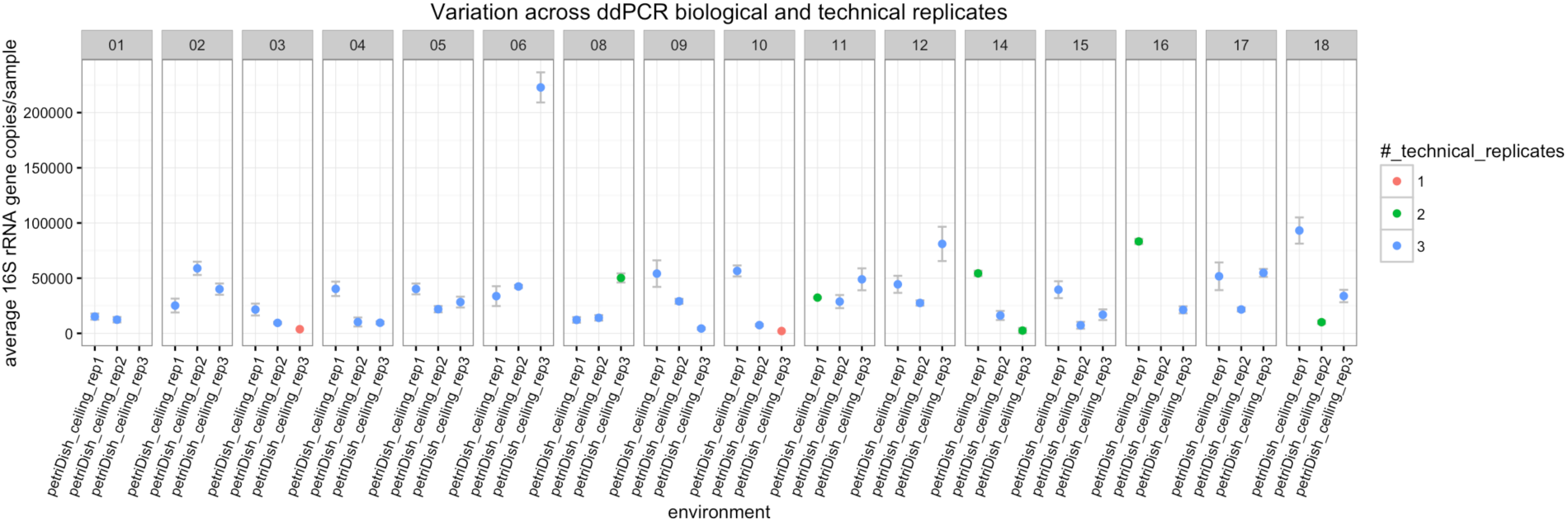
Biological and technical variation across ddPCR replicates. 16S rRNA template copy number was quantified via ddPCR for three petri dish dust collectors suspended from the drop ceiling in each infant’s room. Each dot reflects the average across triplicates runs. Each infant set is labeled at the top of the plot facets.

**Additional file 3:**
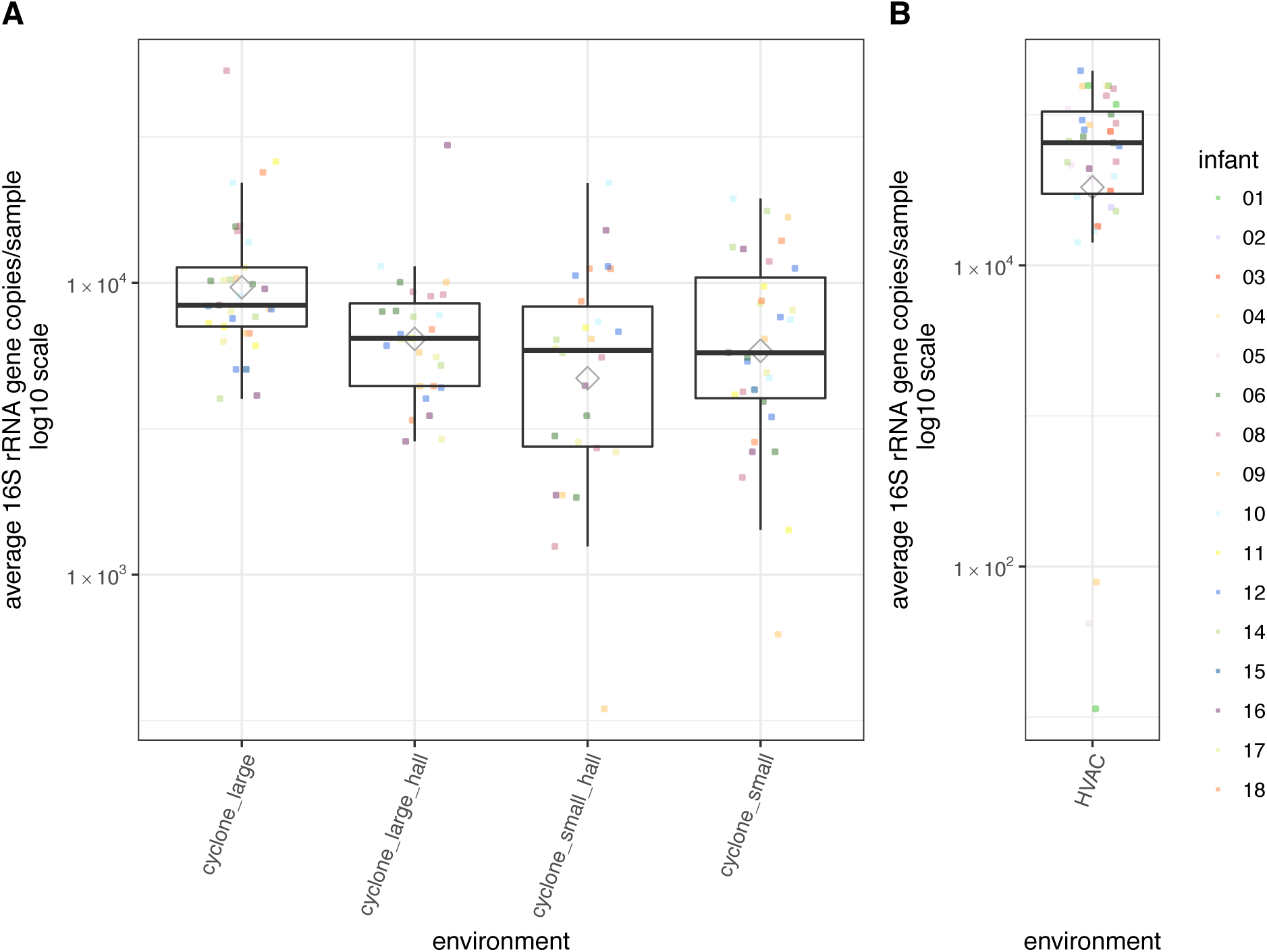
Biomass in air samples from a NICU. 16S rRNA template copy number was quantified via ddPCR. Each dot reflects the average across triplicates runs. Grey diamonds represent averages per environment. Bioaerosol measurements in (A) are separated by small and large size fractions (particles 1-4 μm and > 4 μm, respectively). HVAC samples in (B) were collected from the exterior facet of the HVAC system and represent pretreated air. Counts are normalized per sample per day of collection.

**Additional file 4:**
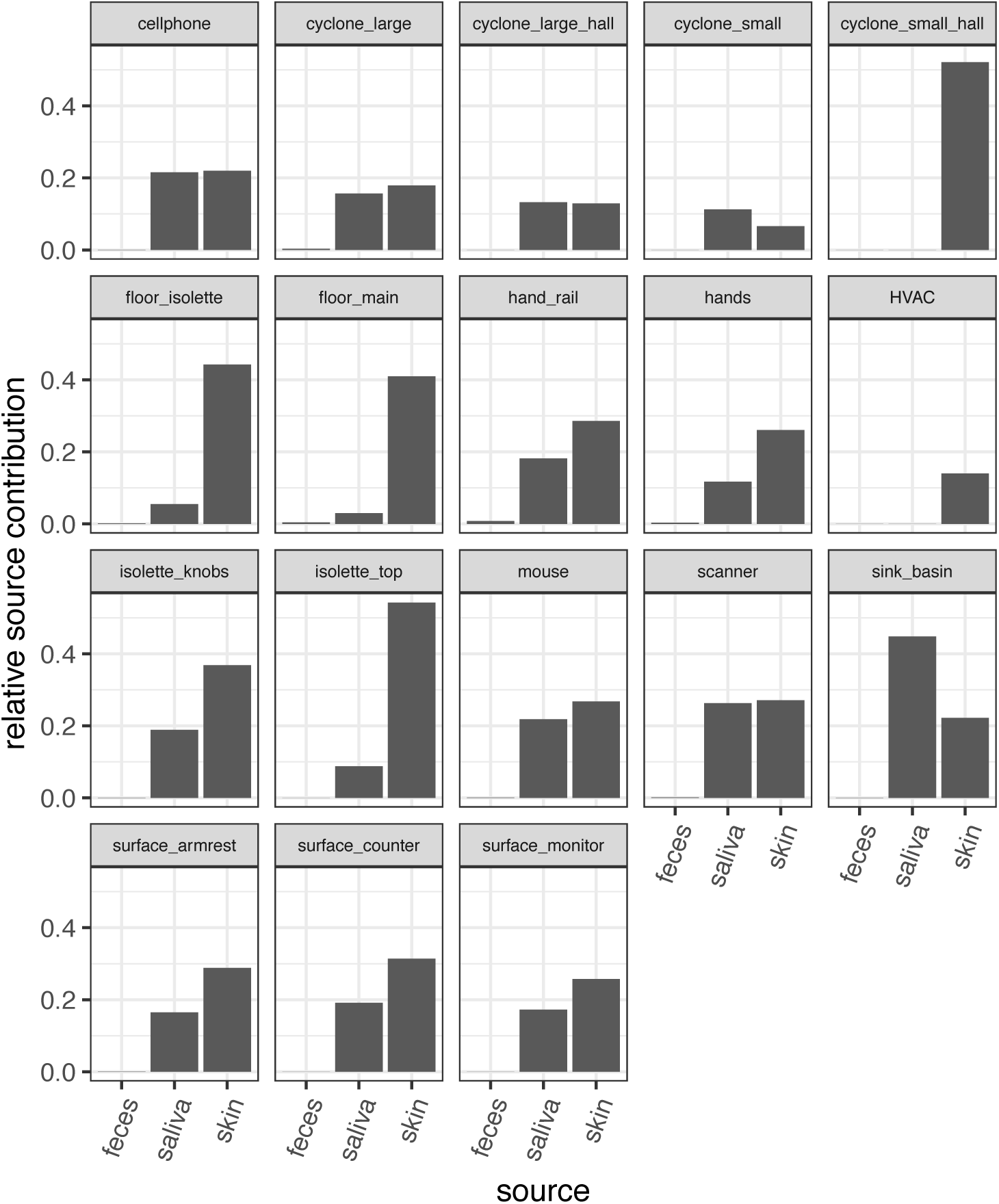
SourceTracker reveals human skin is dominant source of NICU microbes. American Gut skin, oral, and fecal samples were used as “sources” and NICU room samples were used as “sinks” and input into the SourceTracker software. Plotted on the *y*-axis is the mean relative contribution of each human-associated source to each environmental sample.

**Additional file 5:**
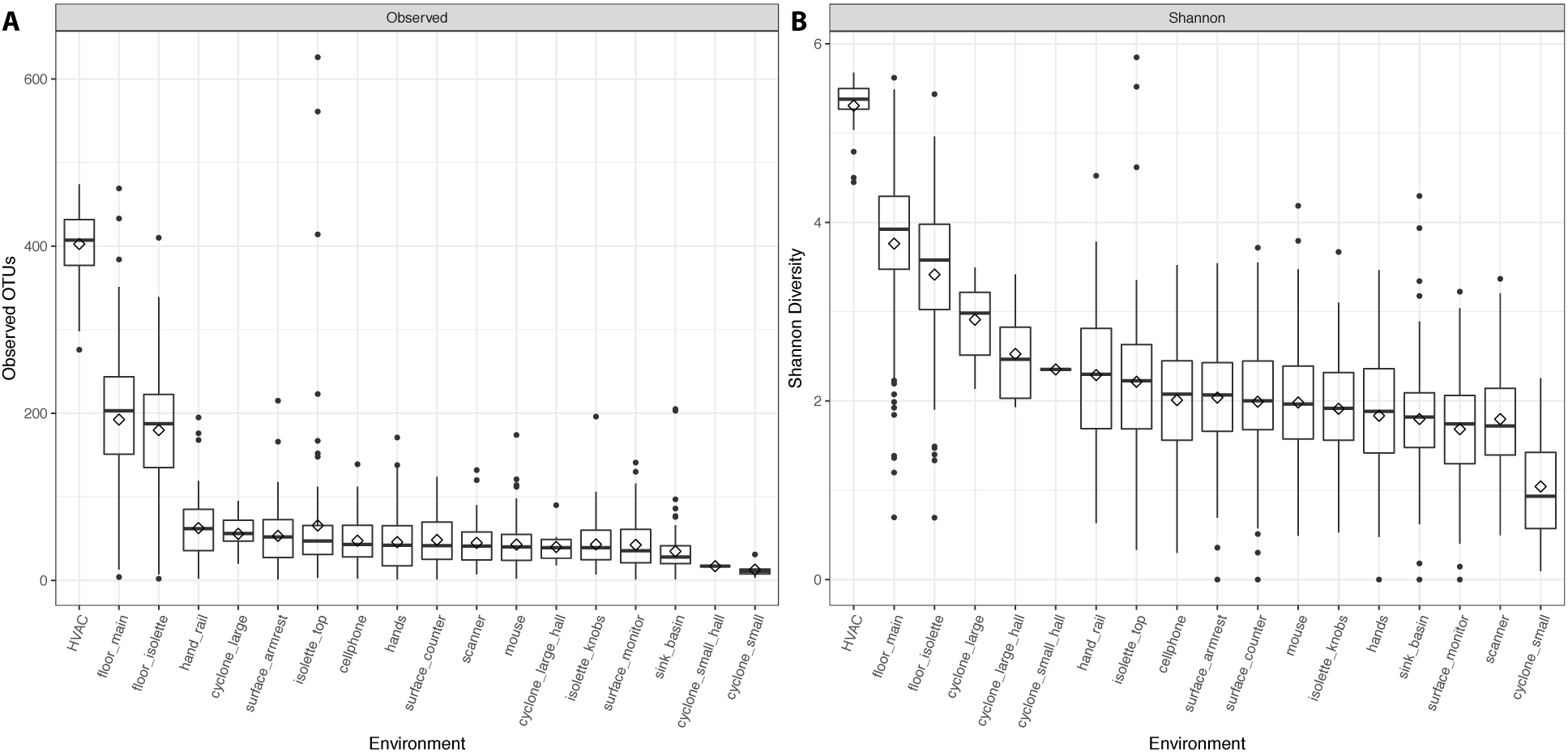
Alpha diversity in a NICU. 16S rRNA amplicon data was used to calculate number of OTUs per environment (a) and the Shannon diversity (b).

**Additional file 6:**
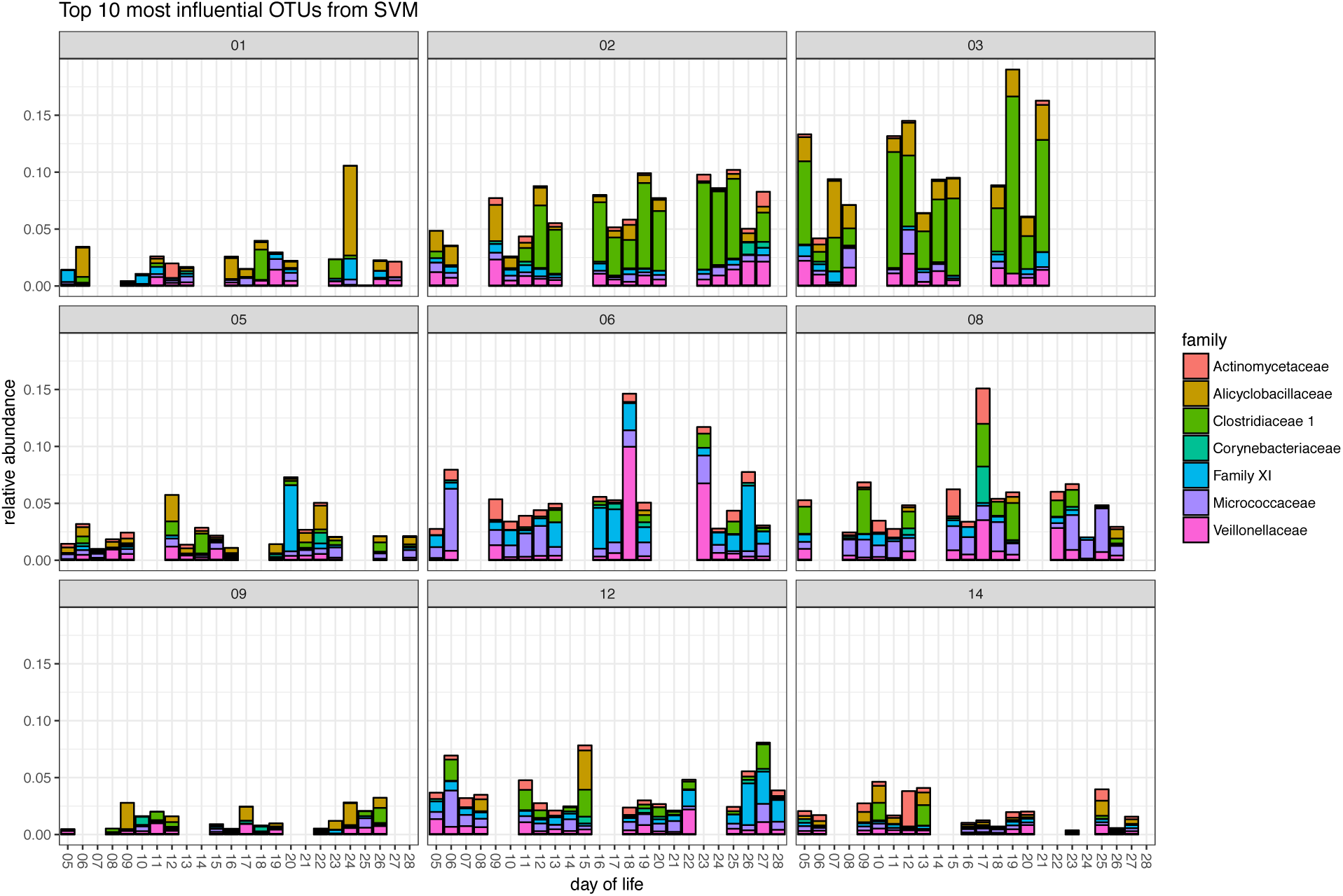
Top 10 most important taxa driving the machine learning model. The top 10 most important variables driving the SVM model are plotted for each infant. On the *y*-axis, “Abundance”, notes the relative importance.

**Additional file 7:**
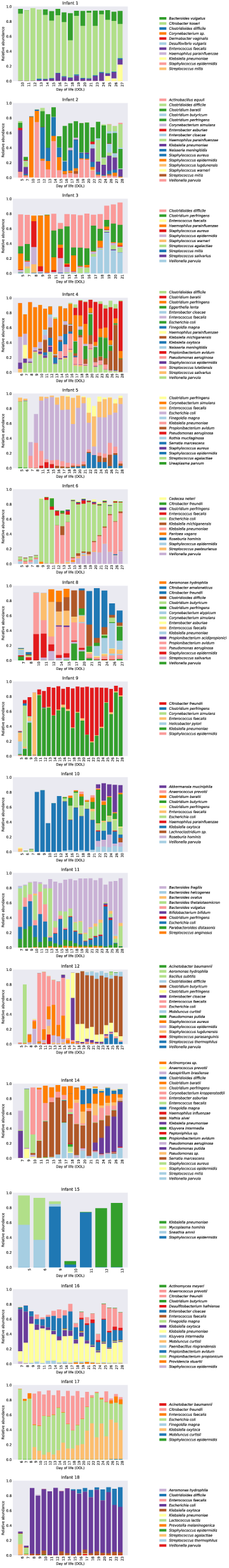
Fecal sample community composition. Plotted in each panel is the community composition of each infant’s fecal samples derived from metagenomics data.

**Additional file 8:**
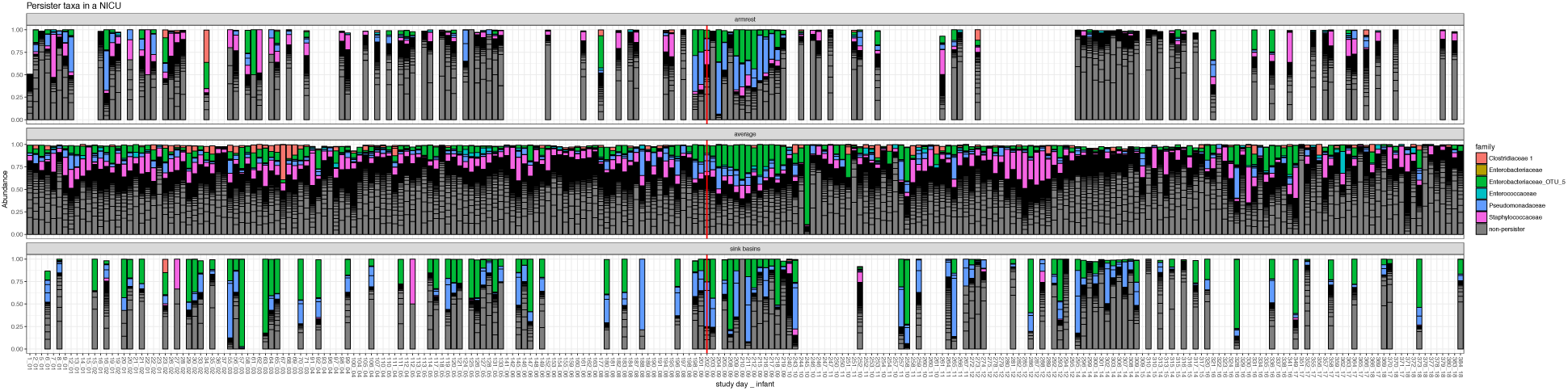
Episodic increases in persistent taxa. The “average” panel represents 16S amplicon data averaged across all samples at each time point per infant. The “armrest” and “sink_basins” panel is the same data but without averaging across environments. The red line highlights the time point in which an increase of *Enterobacteriaceae* was detected in infant 9’s gut. Samples are plotted in chronological order on the x-axis. The plot is split across two pages for clarity.

## References

1. Healthcare-associated Infections [http://www.cdc.gov/winnablebattles/healthcareassociatedinfections/]

2. Arrieta M-C, Stiemsma LT, Dimitriu PA, Thorson L, Russell S, Yurist-Doutsch S, Kuzeljevic B, Gold MJ, Britton HM, Lefebvre DL, Subbarao P, Mandhane P, Becker A, McNagny KM, Sears MR, Kollmann T, Mohn WW, Turvey SE, Brett Finlay B: Early infancy microbial and metabolic alterations affect risk of childhood asthma. Sci Transl Med 2015, 7:307ra152.

3. Cahenzli J, Köller Y, Wyss M, Geuking MB, McCoy KD: Intestinal microbial diversity during early-life colonization shapes long-term IgE levels. Cell Host Microbe 2013, 14:559–570.

4. Gasparrini AJ, Crofts TS, Gibson MK, Tarr PI, Warner BB, Dantas G: Antibiotic perturbation of the preterm infant gut microbiome and resistome. Gut Microbes 2016, 7:443–9.

5. Raveh-Sadka T, Thomas BC, Singh A, Firek B, Brooks B, Castelle CJ, Sharon I, Baker R, Good M, Morowitz MJ, Banfield JF: Gut bacteria are rarely shared by co-hospitalized premature infants, regardless of necrotizing enterocolitis development. Elife 2015, 2015:1– 25.

6. Groer MW, Luciano AA, Dishaw LJ, Ashmeade TL, Miller E, Gilbert JA: Development of the preterm infant gut microbiome: a research priority. Microbiome 2014, 2:38.

7. Hu H, Johani K, Gosbell IB, Jacombs ASW, Almatroudi A, Whiteley GS, Deva AK, Jensen S, Vickery K: Intensive care unit environmental surfaces are contaminated by multidrug-resistant bacteria in biofilms: Combined results of conventional culture, pyrosequencing, scanning electron microscopy, and confocal laser microscopy. J Hosp Infect 2015, 91:35–44.

8. Shin H, Pei Z, Martinez KA, Rivera-Vinas JI, Mendez K, Cavallin H, Dominguez-Bello MG: The first microbial environment of infants born by C-section: the operating room microbes. Microbiome 2015, 3:59.

9. Bäckhed F, Roswall J, Peng Y, Feng Q, Jia H, Kovatcheva-Datchary P, Li Y, Xia Y, Xie H, Zhong H, Khan MT, Zhang J, Li J, Xiao L, Al-Aama J, Zhang D, Lee YS, Kotowska D, Colding C, Tremaroli V, Yin Y, Bergman S, Xu X, Madsen L, Kristiansen K, Dahlgren J, Jun W: Dynamics and stabilization of the human gut microbiome during the first year of life. Cell Host Microbe 2015, 17:690–703.

10. Chu DM, Ma J, Prince AL, Antony KM, Seferovic MD, Aagaard KM: Maturation of the infant microbiome community structure and function across multiple body sites and in relation to mode of delivery. Nat Publ Gr 2017(August 2016).

11. Dominguez-Bello MG, Costello EK, Contreras M, Magris M, Hidalgo G, Fierer N, Knight R: Delivery mode shapes the acquisition and structure of the initial microbiota across multiple body habitats in newborns. Proc Natl Acad Sci U S A 2010, 107:11971–5.

12. Brooks B, Firek BBA, Miller CCS, Sharon I, Thomas BC, Baker R, Morowitz MJ, Banfield JF: Microbes in the neonatal intensive care unit resemble those found in the gut of premature infants. Microbiome 2014, 2:1.

13. Miller CS, Baker BJ, Thomas BC, Singer SW, Banfield JF: EMIRGE: reconstruction of full-length ribosomal genes from microbial community short read sequencing data. Genome Biol 2011, 12:R44.

14. Raveh-Sadka T, Firek B, Sharon I, Baker R, Brown CT, Thomas BC, Morowitz MJ, Banfield JF: Evidence for persistent and shared bacterial strains against a background of largely unique gut colonization in hospitalized premature infants. ISME J 2016, 10:2817–2830.

15. Gibson MK, Wang B, Ahmadi S, Burnham C-AD, Tarr PI, Warner BB, Dantas G: Developmental dynamics of the preterm infant gut microbiota and antibiotic resistome. Nat Microbiol 2016, 1:16024.

16. Buffet-Bataillon S, Branger B, Cormier M, Bonnaure-Mallet M, Jolivet-Gougeon A: Effect of higher minimum inhibitory concentrations of quaternary ammonium compounds in clinical *E. coli* isolates on antibiotic susceptibilities and clinical outcomes. J Hosp Infect 2011, 79:141–6.

17. Strain-resolved community genomic analysis of gut microbial colonization in a premature infant Michael J. Morowitz. 2010.

18. Licina D, Bhangar S, Brooks B, Baker R, Firek B, Tang X, Morowitz MJ, Banfield JF, Nazaroff WW, Berkeley UC, Jillian F, William W: Concentrations and sources of airborne particles in a neonatal intensive care unit. PLoS One 2016, 11:e0154991.

19. Lindsley WG, Blachere FM, Thewlis RE, Vishnu A, Davis KA, Cao G, Palmer JE, Clark KE, Fisher MA, Khakoo R, Beezhold DH: Measurements of airborne influenza virus in aerosol particles from human coughs. PLoS One 2010, 5:e15100.

20. Adams RI, Miletto M, Taylor JW, Bruns TD: Dispersal in microbes: fungi in indoor air are dominated by outdoor air and show dispersal limitation at short distances. ISME J 2013, 7:1262–73.

21. Yamamoto N, Shendell DG, Peccia J: Assessing allergenic fungi in house dust by floor wipe sampling and quantitative PCR. Indoor Air 2011, 21:521–30.

22. Fadrosh DW, Ma B, Gajer P, Sengamalay N, Ott S, Brotman RM, Ravel J: An improved dual-indexing approach for multiplexed 16S rRNA gene sequencing on the Illumina MiSeq platform. Microbiome 2014, 2:6.

23. Hildebrand F, Tadeo R, Voigt AY, Bork P, Raes J: LotuS: an efficient and user-friendly OTU processing pipeline. Microbiome 2014, 2:1–7.

24. Brooks B, Olm MR, Firek BA, Baker R, Thomas BC, Morowitz MJ, Banfield JF: Strain-resolved analysis of hospital rooms and infants reveals overlap between the human and room microbiome. Nat Commun 2017, In Review.

25. Lazarevic V, Gaïa N, Girard M, Schrenzel J: Decontamination of 16S rRNA gene amplicon sequence datasets based on bacterial load assessment by qPCR. BMC Microbiol 2016, 16:73.

26. Meadow JF, Altrichter AE, Bateman AC, Stenson J, Brown G, Green JL, Bohannan BJM: Humans differ in their personal microbial cloud. PeerJ 2015, 3:e1258.

27. Knights D, Kuczynski J, Charlson E, Zaneveld J, Mozer MC, Collman RG, Bushman FD, Knight R, Kelley ST: Bayesian community-wide culture-independent microbial source tracking. Nat Methods 2011, 8:761–3.

28. Chase J, Fouquier J, Zare M, Sonderegger DL, Knight R, Kelley ST, Siegel J, Caporaso JG: Geography and location are the primary drivers of office microbiome composition. mSystems 2016, 1:e00022–16.

29. Jovel J, Patterson J, Wang W, Hotte N, O’Keefe S, Mitchel T, Perry T, Kao D, Mason AL, Madsen KL, Wong GKS: Characterization of the gut microbiome using 16S or shotgun metagenomics. Front Microbiol 2016, 7(APR):1–17.

30. Edgar RC: Search and clustering orders of magnitude faster than BLAST. Bioinformatics 2010, 26:2460–1.

31. Yuan C, Lei J, Cole J, Sun Y: Reconstructing 16S rRNA genes in metagenomic data. Bioinformatics 2015, 31:i35–i43.

32. Luoma M, Batterman SA: Characterization of particulate emissions from occupant activities in offices. Indoor Air 2001, 11:35–48.

33. Fairchild CI, Tillery MI: Wind tunnel measurements of the resuspension of ideal particles. Atmos Environ 1982, 16:229–38.

34. Bhangar S, Brooks B, Firek B, Licina D, Tang X, Morowitz MJ, Banfield JF, Nazaroff WW: Pilot study of sources and concentrations of size-resolved airborne particles in a neonatal intensive care unit. Build Environ 2016, 106:10–19.

35. Kembel SW, Jones E, Kline J, Northcutt D, Stenson J, Womack AM, Bohannan BJ, Brown GZ, Green JL: Architectural design influences the diversity and structure of the built environment microbiome. ISME J 2012, 6:1469–79.

36. Kembel SW, Meadow JF, O’Connor TK, Mhuireach G, Northcutt D, Kline J, Moriyama M, Brown GZ, Bohannan BJM, Green JL: Architectural design drives the biogeography of indoor bacterial communities. PLoS One 2014, 9:e87093.

37. Luangasanatip N, Hongsuwan M, Limmathurotsakul D, Lubell Y, Lee AS, Harbarth S, Day NPJ, Graves N, Cooper BS: Comparative efficacy of interventions to promote hand hygiene in hospital: systematic review and network meta-analysis. BMJ 2015, 351:h3728.

38. Lax S, Sangwan N, Smith D, Larsen P, Handley KM, Richardson M, Guyton K, Krezalek M, Shogan BD, Defazio J, Flemming I, Shakhsheer B, Weber S, Landon E, Garcia-Houchins S, Siegel J, Alverdy J, Knight R, Stephens B, Gilbert JA: Bacterial colonization and succession in a newly opened hospital. Sci Transl Med 2017, 9:1–11.

39. Romanova NA, Gawande P V, Brovko LY, Griffiths MW: Rapid methods to assess sanitizing efficacy of benzalkonium chloride to *Listeria monocytogenes* biofilms. J Microbiol Methods 2007, 71:231–7.

40. Weiss-Muszkat M, Shakh D, Zhou Y, Pinto R, Belausov E, Chapman MR, Sela S: Biofilm formation by and multicellular behavior of *Escherichia coli* O55:H7, an atypical enteropathogenic strain. Appl Environ Microbiol 2010, 76:1545–54.

41. Hoffman LR, D’Argenio DA, MacCoss MJ, Zhang Z, Jones RA, Miller SI: Aminoglycoside antibiotics induce bacterial biofilm formation. Nature 2005, 436:1171–5.

42. Bokulich NA, Mills DA, Underwood M a.: Surface microbes in the neonatal intensive care unit: Changes with routine cleaning and over time. J Clin Microbiol 2013, 51:2617–24.

